# A curriculum learning approach to training antibody language models

**DOI:** 10.1101/2025.02.27.640641

**Authors:** Sarah M. Burbach, Bryan Briney

## Abstract

There is growing interest in pre-training antibody language models (**AbLMs**) with a mixture of unpaired and natively paired sequences, seeking to combine the proven benefits of training with natively paired sequences with the massive scale of unpaired antibody sequence datasets. However, given the novelty of this strategy, the field lacks a systematic evaluation of data processing methods and training strategies that maximize the benefits of mixed training data while accommodating the significant imbalance in the size of existing paired and unpaired datasets. Here we introduce a method of curriculum learning for AbLMs, which facilitates a gradual transition from unpaired to paired sequences during training. We optimize this method and show that a 650M-parameter curriculum model, CurrAb, outperforms existing mixed AbLMs in downstream classification tasks.

## INTRODUCTION

Antibodies are a diverse and essential component of the adaptive immune system, with total available repertoire diversity estimated as high as 10^18^ unique antibodies.^1^ This exceptional diversity results initially from the somatic recombination of germline gene segments and is further refined upon antigen exposure via clonal expansion, somatic hypermutation (**SHM**), and antigen-driven selection of productive mutations. Within each recombined antibody gene, diversity is greatest in the complementary determining regions (**CDRs**), which comprise the antigen-recognition site and thus determine antibody specificity.

Given the enormous diversity of the antibody repertoire, antibody language models (**AbLMs**) have grown in popularity for their potential to speed up the discovery and engineering of novel antibodies, particularly due to their ability to learn immunological mechanisms such as affinity maturation.^2^ Because antibody recombination is a modular process, AbLMs tend to learn germline-encoded features easily but struggle with mutated residues and in the primarily non-templated CDR3.^3,4^ Given this fact, a major focus of current studies is to improve model performance in the CDR regions. We have recently shown that pre-training an AbLM with natively paired antibody sequences rather than unpaired sequences improves the model’s ability to learn immunologically significant features that span both heavy and light chains, including heightened cross-chain attention on the functionally critical CDRs.^4^

Despite the proven training benefits of paired sequences, available paired antibody sequence datasets are much smaller than unpaired datasets, and it is well documented that the cross-entropy (**CE**) loss of language models (**LMs**) scales with dataset size.^5,6^ Given this, recent models^3,7,8^ have incorporated a mix of unpaired and natively paired sequences, with the intuition that we can supplement the limited paired data with the larger quantities of unpaired data available. In theory, this should also improve a model’s ability to learn non-germline residues, since it sees a more diverse collection of somatically mutated antibodies in the larger unpaired datasets. Models trained with a mixture of unpaired and paired data typically pre-train using unpaired sequences and fine-tune with paired sequences.^3^ IgBert and IgT5 from Kenlay et al 2024^7^ introduced a variation of this approach, using a 2:1 ratio of unpaired to paired data during the fine-tuning phase to help mitigate catastrophic forgetting.^9^

However, due to the recency of this pre-training shift, we lack a systematic evaluation of pretraining strategies to determine the optimal parameters for a model trained on a mixture of these datatypes. We theorized that a smoother transition between unpaired and paired sequences may help mitigate the risk of catastrophic forgetting further. Specifically, we introduce a method of training AbLMs inspired by curriculum learning,^10^ which has been successful in the natural language processing domain. Curriculum learning involves ordering the training data, traditionally from ‘easiest’ to ‘hardest’. One such method is length-based curriculum learning, where training begins with shorter examples and transitions to longer ones.^11^ To apply this to AbLMs, we ordered the data to start with the less information-dense unpaired sequences and then transition to the more information-dense paired sequences gradually during training. We hypothesized that this may enable the model to balance its knowledge of both sequence types without catastrophic forgetting while still emphasizing the paired sequences. We will compare this curriculum approach with other training methods including single data-type models, the traditional fine-tuning approach, and a constant approach (where data is mixed throughout training). In addition, we will train a large 650M-parameter curriculum model, CurrAb, and compare its performance on downstream tasks to existing mixed AbLMs.

## RESULTS

### Implementing curriculum learning for antibodies

Given the unique challenges associated with pre-training on both unpaired and paired sequence data, we introduce a modified version of curriculum learning for AbLMs. This method modifies the way training examples are sampled, such that a variable percentage of unpaired sequences is selected based on the training step. The highest percentage of unpaired sequences are selected at the beginning of training and this percentage decreases as training progresses, to emphasize paired sequences in the final stages of training. To implement this, we generate an unpaired probability curve that determines what fraction of each training batch should be sampled from unpaired and paired datasets at each step of training (***Fig 1***). The probability curve is a sigmoid decay function that enables the gradual transition from unpaired to paired sequences that can be modified to match training needs.

**Figure 1.**
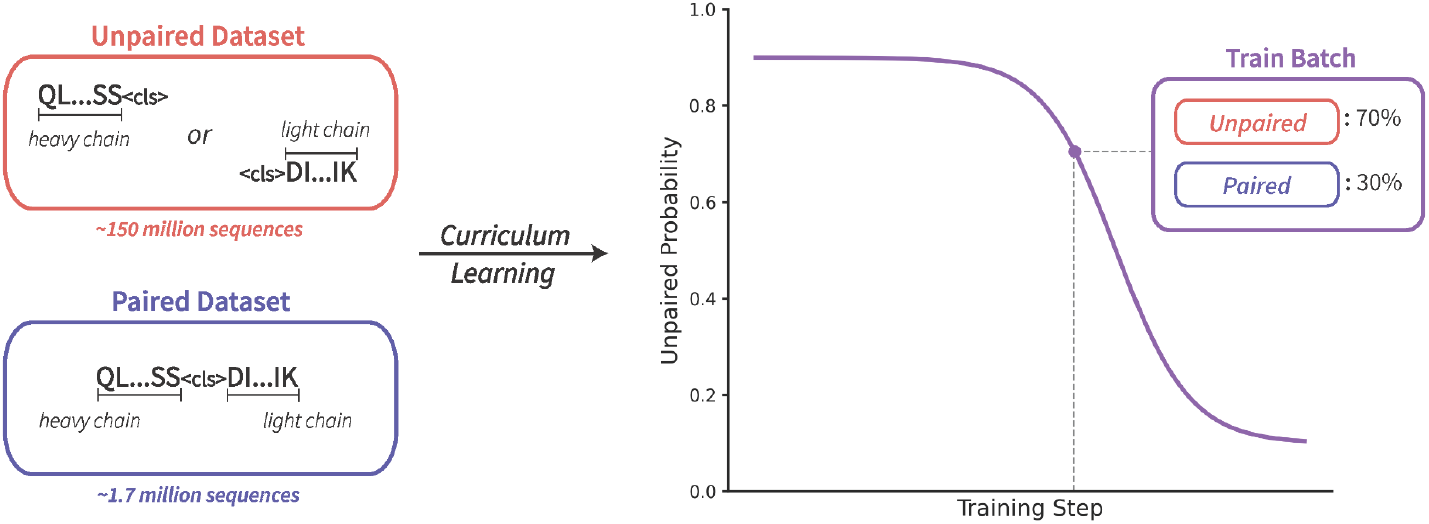
Curriculum learning implementation for AbLMs. The approach is designed to help handle the massive data imbalance between unpaired and paired datasets. To implement this, an unpaired probability curve determines the percentage of data sampled from the unpaired dataset for a given batch at a given step. This percentage decreases as training progresses, to allow for an emphasis on paired sequences near the end of training.

To optimize the curriculum strategy, we tested a variety of hyperparameters general to mixed-data models as well as curriculum-specific parameters, such as modifications to the unpaired probability curve and the learning rate schedule. To efficiently evaluate a large number of pre-training parameters, we initially used a pilot-scale 55M parameter variant of the ESM-2^12^ architecture, with a slightly modified vocab of 33 characters that includes a unique <sep> token. These models were trained for 100k steps on a subset (20%) of the full paired and unpaired datasets.

### Optimizing model architecture and data processing for mixed-data models

Prior to implementing the curriculum learning approach, we assessed the impact of a few key architecture and data formatting parameters for mixed-data models generally. Most significant are the changes observed based on the positional embedding (**PE**) type. Rotary position embeddings (**RoPE**)^13^ consistently allow LMs to converge more quickly and reach a lower final loss than absolute PEs. By rotating the keys and queries based on their relative distance, RoPE can better accommodate sequences of varying lengths. Given that paired antibody sequences are approximately twice the length of unpaired ones, we hypothesized that RoPE would be especially impactful for AbLMs trained with a mixture of paired and unpaired sequences. To test this, we trained four models: two trained using only paired sequences (paired-only models) and two trained using only unpaired sequences (unpaired-only models), with either RoPE or absolute PE. Each of these models was then evaluated using both paired and unpaired test datasets. As expected, RoPE models perform slightly better on cross-entropy (**CE**) loss than absolute PE models when tested with the same data type they were trained on (paired-only models tested with paired sequences and unpaired-only models tested with unpaired sequences). Notably, when evaluated using heterologous data (paired test sequences for unpaired-only models or unpaired test sequences for paired-only models), RoPE models outperform absolute PE models by a large margin (***Fig 2A, 2C***). In the paired-only models, the absolute PE model performs similarly to the RoPE model in tests containing unpaired heavy chains, but significantly worse across all regions in unpaired light chains (***Fig 2B***). Similarly, in the unpaired-only models, the performance of the absolute PE model on paired sequences gets progressively worse in the heavy chain and is significantly worse in the light chain, compared to the RoPE model (***Fig 2D***). Our observation that the majority of the difference in performance between absolute PE and RoPE models is observed in the light chains, may be explained by their placement after heavy chains in paired sequence inputs.

**Figure 2.**
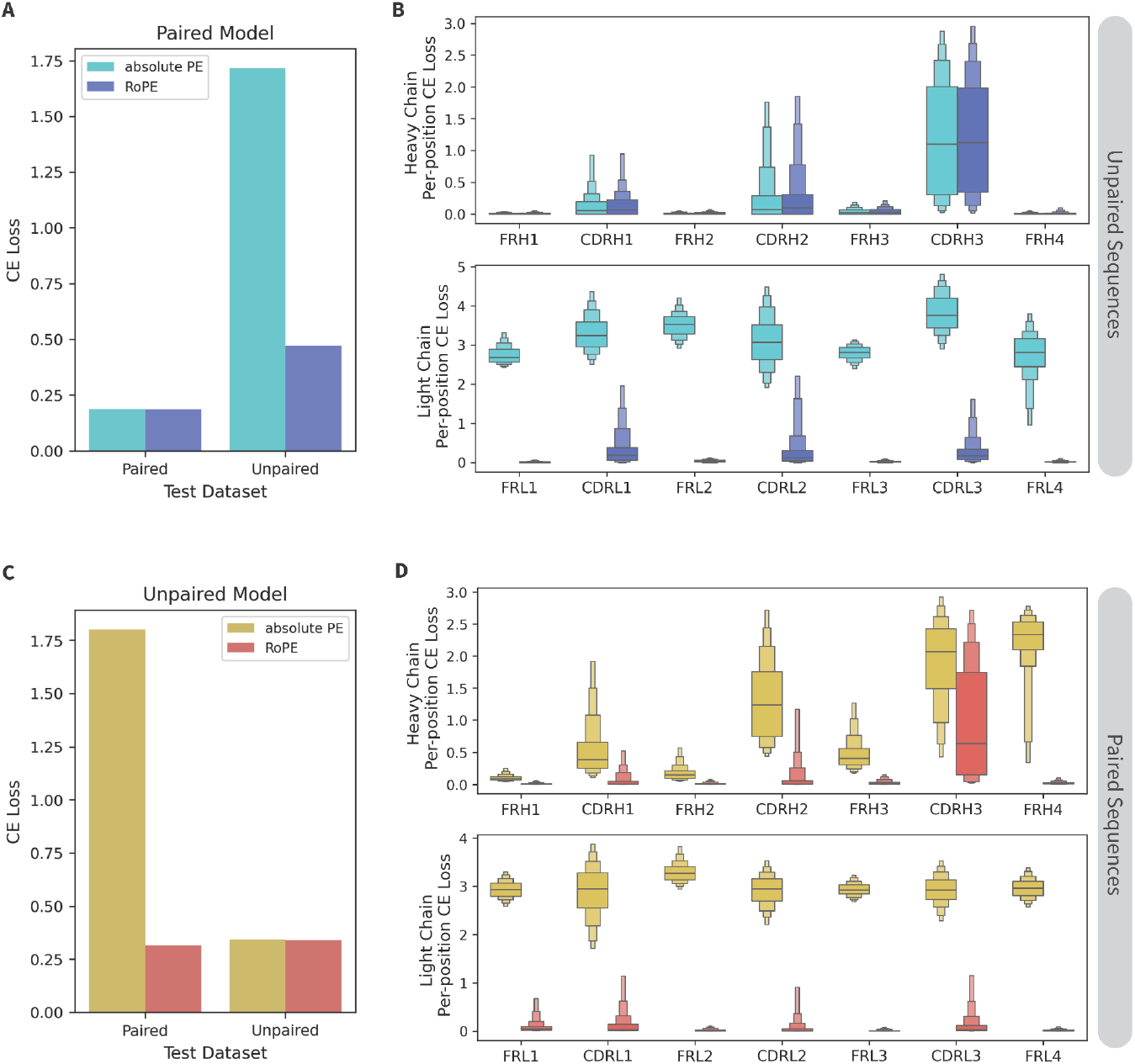
Comparing RoPE and absolute PE with paired and unpaired models. Two (A-B) paired-only model and (C-D) two unpaired-only models were trained with either RoPE or absolute PE. (A, C) CE loss on paired and unpaired test datasets of ∼10k sequences each. (B) Per-position CE loss of the paired-only models on 1k sequences from the unpaired test dataset. (C) Per-position CE loss of the unpaired-only models on 1k sequences from the paired test dataset.

We next tested different separator token(s) and varying ratios of unpaired:paired data in mixed-data models. First, we note that a single separator token produced slight but consistent improvement on both CE loss and classifier performance, regardless of whether the separator token was uniquely used for chain separation (<sep>) or a reuse of the start-of-string token (<cls>) (***Table S1***). Interestingly, adding a separator to unpaired sequences (immediately following heavy chains or immediately preceding light chains) resulted in better performance than omitting the separator, suggesting that a separator is useful even in an unpaired-only model. Second, we observe that increasing training time for one data type consistently results in a lower loss of that data type, but it always comes at the cost of increased loss on the other data type. Once the total training dataset exceeds 50% paired sequences, the paired loss fails to improve any further (***Table S2***). Based on this, we train all subsequent mixed-data models with 62.5% unpaired data (just over 1 epoch of our total unpaired training dataset).

### Optimizing curriculum learning for antibodies

To optimize our curriculum learning method, we tested a variety of modifications to the unpaired probability curve and learning rate (**LR**) schedule. We first tested different ranges of probabilities for these curves, with the widest ranging from 1.0 to 0.0 (max1) and the narrowest ranging from 0.7 to 0.3 (max0.7) (***Fig 3A***). Performance on the unpaired test set reveals that the CE loss decreases as the probability range narrows (***Fig 3B***). Performance on the paired test set is much less variable, with the max0.8 range resulting in the lowest loss by a small margin. These results suggest that including a mix of data types (with a minimum of ∼20% of the minority data type) throughout training is useful to the models’ final performance. Second, we tested the impact of the slope of the unpaired probability curve decay, which is determined by the k-value. Models with a larger k-value decay at a steeper rate than models with a smaller k-value, resulting in a more rapid transition from unpaired to paired training data (***Fig 3C***). Changing the slope has almost no effect on the paired performance, with k = 50 performing only slightly better than the others (***Fig 3D***). We observe a slightly larger impact on the unpaired loss, with k = 15 resulting in the lowest CE loss.

**Figure 3.**
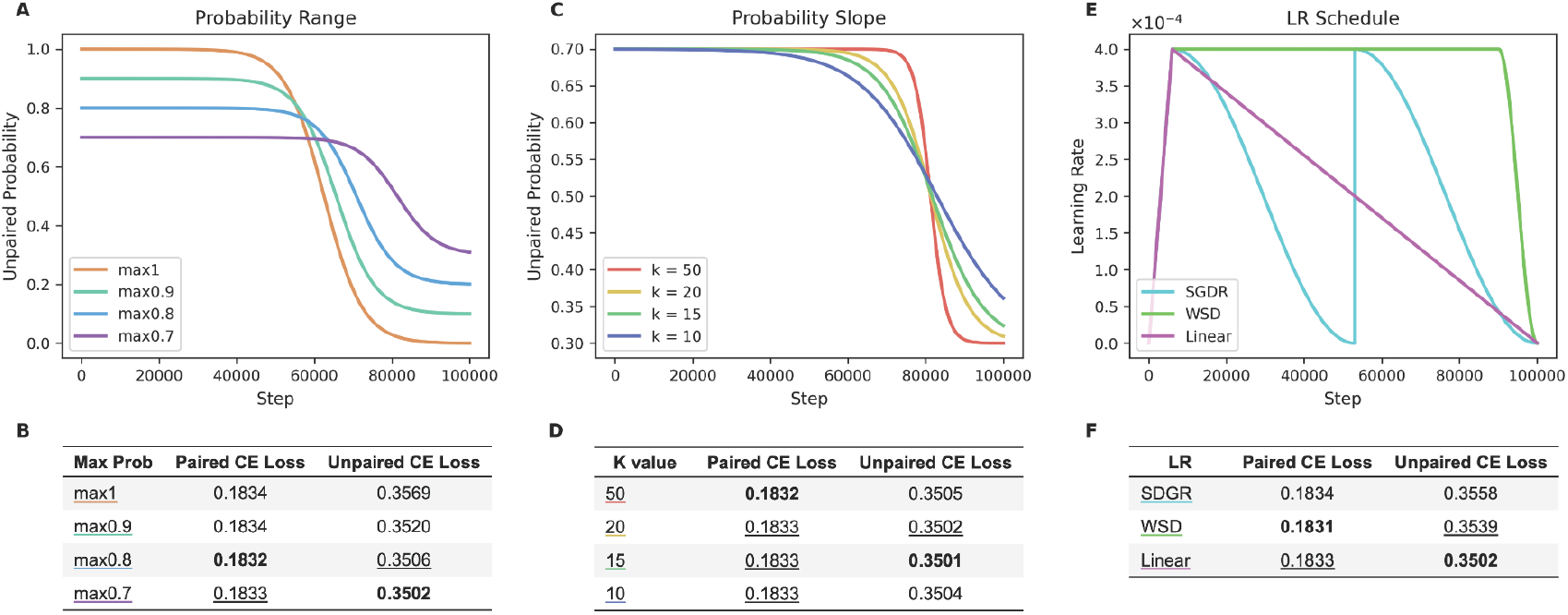
Optimizing hyperparameters for curriculum models. (A-B) Testing different ranges of the unpaired probability curve. (C-D) Testing different slopes of the unpaired probability curve, by adjusting their k-value. (E-F) Testing three different LR schedules: linear, WSD, and SGDR. All models were evaluated based on their CE loss on the paired and unpaired test datasets.

Finally, we tested different learning rate schedules to determine if a higher learning rate later in training is useful for learning paired sequences. We evaluated a linear-decay schedule, a warmup-stable-decay (**WSD**)^14^ schedule, and a cosine annealing schedule (commonly known as stochastic gradient descent with warm restarts (**SGDR**))^15^ (***Fig 3E***). We find that the linear LR results in the lowest unpaired test loss, while WSD results in the lowest test loss for paired sequences (***Fig 3F***). The improved performance of the WSD LR suggests that a higher learning rate during the transition can assist the model in learning paired sequences. However, this small gain in paired performance comes at a much larger cost to unpaired performance.

Based on the limited improvements observed in the paired CE loss, we optimized the curriculum implementation based on the unpaired performance. Therefore, our optimized parameters are a probability range of 0.7 to 0.3, a k-value of 15, and a linear LR schedule.

### Assessing methods for training mixed models

To directly compare strategies for training with mixed datasets, we trained a series of five models: one model pre-trained on unpaired data and finetuned on paired data (*finetuned*); one model trained with a constant ratio of paired and unpaired data (*constant*), one model trained using our curriculum-training strategy that dynamically adjusts the ratio of paired and unpaired data throughout training (*curriculum*), and two control models trained using only unpaired (*unpaired-only*) or paired (*paired-only*) data (***Fig 4A***). Each model was otherwise identical in architecture and was trained for 100k steps. The 50k checkpoint was used for downstream testing of the paired-only model due to overfitting. The paired:unpaired ratio of the constant model and finetuning duration of the finetuned model were selected to ensure that all mixed models were trained with the same total amount (in epochs) of unpaired and paired training data.

**Figure 4.**
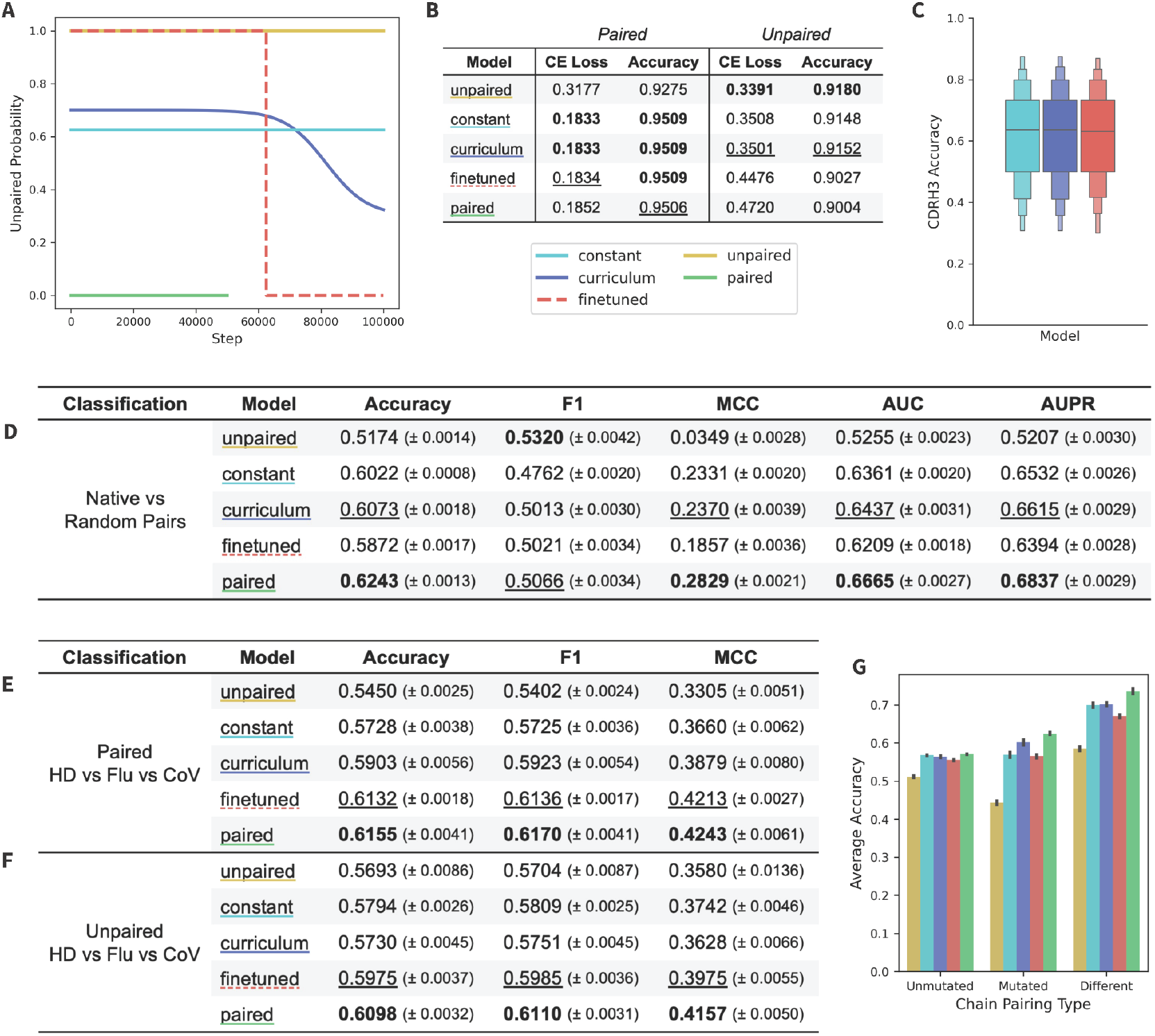
Comparing the performance of mixed model training methods. (A) Unpaired probability curves for the five models. (B) CE loss on paired and unpaired test datasets of ∼10k sequences. (C) Mixed models accuracy at predicting CDRH3 of 1k paired sequences from the test set. Results for the three classification tasks, which are one (D) *Native vs Random* pair classification and two *Healthy Donor vs Flu vs CoV* specificity classifications using (E) paired and (F) unpaired sequences. (G) The results of the pair classification were additionally split by mutated pairs, unmutated pairs, and mismatched pairs. Metrics on classification tasks are mean and standard error, with the highest values bolded and the second highest values underlined.

We evaluated each model with a masked language modeling objective using the held-out test data (***Fig 4B***) and observed that the curriculum and constant models performed best across CE loss and accuracy on the paired test set. We additionally observed that the unpaired-only model performed best across both CE loss and accuracy on the unpaired test set, followed by the curriculum model. The boost in unpaired performance in the curriculum model compared to the other mixed models suggests that the curriculum training schedule enables the model to retain its knowledge of unpaired sequences the best. To further assess the paired accuracy of the mixed models, we also calculated the CDRH3 accuracy for these three models (***Fig 4C***). We observed that the CDRH3 accuracy is essentially the same across all mixed models, indicating that the method of training a mixed model does not have a significant impact on the model’s understanding of the CDRH3 region.

To further assess model performance, we performed a pair classification task, introduced in Ng and Briney 2024^16^. The goal of this binary classification task is to identify whether a given paired sequence is a native or random pairing. The dataset was generated from the paired test dataset and is composed of naive (unmutated) and memory (mutated) sequences. The paired-only model performed best, followed by the curriculum model which outperforms the other mixed data models (***Fig 4D***). Next, we separated the test data for pair classification into three subsets: pairs in which both chains are mutated (*mutated*), pairs in which both chains are unmutated (*unmutated*), or pairs in which one chain is mutated while the other is not (*different*) (***Fig 4G***). All models except the unpaired model perform significantly better on the different subset, indicating that pair classification is driven, at least in par,t by identifying similar levels of mutation across both chains. Performance on the mutated subset is most varied between models, with the paired-only and curriculum models showing the highest classification accuracy.

We next performed three-way specificity classification tasks, in which the model was finetuned to classify between coronavirus (**CoV**) specific antibodies, influenza (**Flu**) specific antibodies, and nonspecific healthy donor (**HD**) antibodies. Models were fine-tuned with either paired sequences (***Fig 4E***) or unpaired heavy chain sequences (***Fig 4F***). The paired-only model outperformed all models across both the paired and unpaired tasks, followed closely by the finetuned model. Results for a two-way specificity classification task (CoV vs HD) can be found in ***Table S3***, where we observe similar results but exhibit smaller margins of difference. This suggests that a strong emphasis on paired sequences (despite the previously observed forgetting of unpaired sequences) is of primary importance for specificity classification tasks.

The higher performance of the paired-only model compared to the mixed models seems to be an artifact of the small size of our pilot-scale models, however. When the same classification tasks are performed with larger 650M-parameter models, the curriculum model outperforms the paired-only model (***Table S4***) across all tasks.

### Large-scale curriculum model outperforms existing mixed models

A distinct advantage of mixed models over paired-only models is the massively increased training data scale, which could allow longer training of larger models before overfitting. To test the scalability of our curriculum method, we trained CurrAb: a 650M-parameter model based on the ESM-2 architecture. The model was trained for 500k steps using the optimized curriculum schedule described above and a reduced learning rate (1e-4) to stabilize training.

We compared the performance of CurrAb to several existing mixed models: IgBERT^7^, IgT5^7^, AbLang2^3^, and AntiBERTa2^8^, using the pair classification task described previously. CurrAb performed better than all other mixed models across all metrics except F1, followed most closely by AbLang2 (***Fig 5A***). The unexpectedly low performance of IgBERT and AntiBERTa2 may be due to the classification head of BERT models, which is only a single projection layer that may not be sufficient for this complex task. We additionally performed the three-way specificity classification with CoV-specific, Flu-specific, and HD sequences. CurrAb again outperformed other mixed models on both the paired (***Fig 5B***) and unpaired (***Fig 5C***) specificity classification tasks. The second-best model performances are AbLang2 in paired specificity classification and AntiBERTa2 in unpaired specificity classification. A closer examination of the paired specificity classification results showed that CurrAb was >10% more accurate than the next best-performing model, AbLang2 (***Fig 5D***).

**Figure 5.**
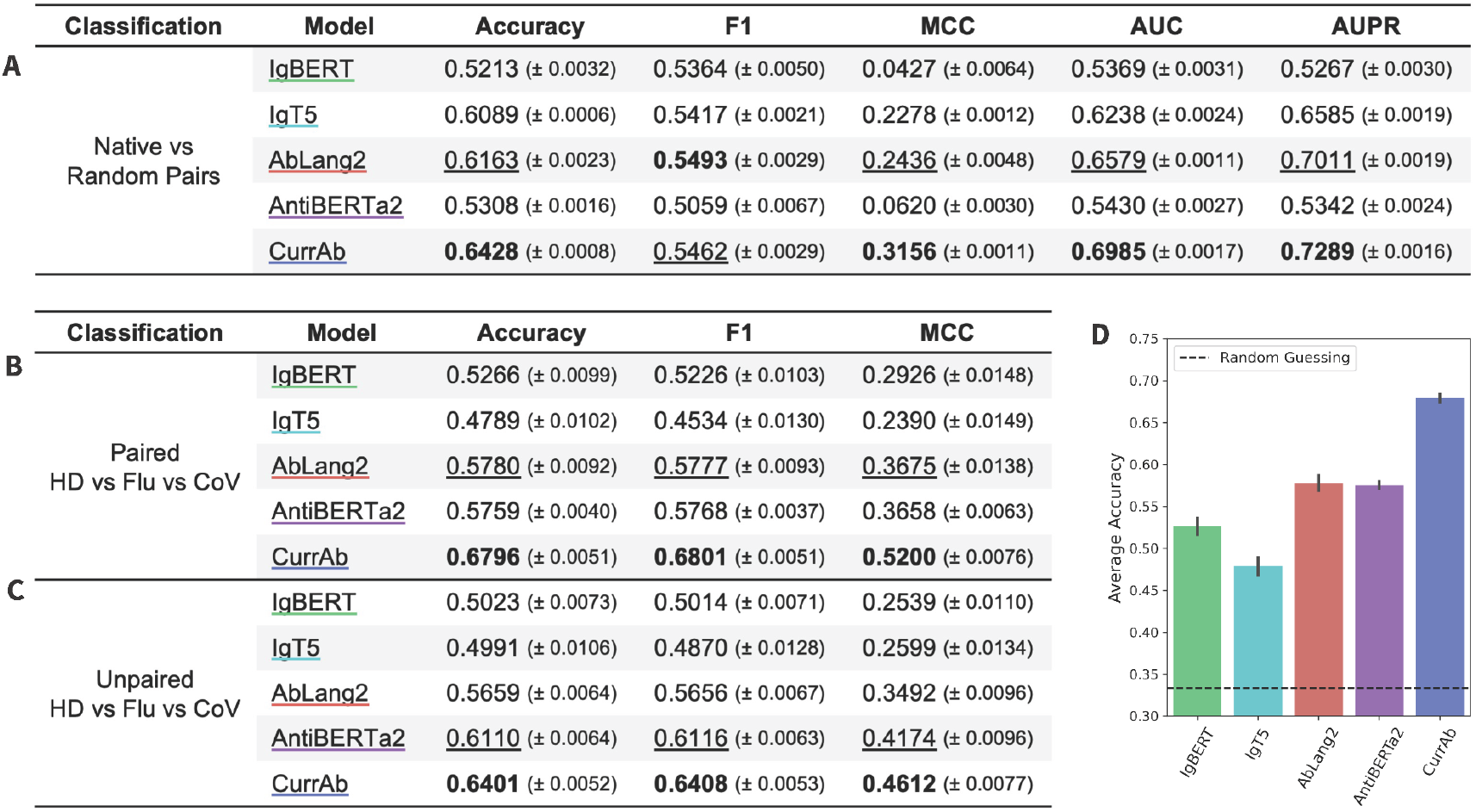
Comparing CurrAb to existing mixed data models. Results of three classification tasks, which are one (A) *Native vs Random* pair classification and two *Healthy Donor vs Flu vs CoV* specificity classification tasks using (B) paired and (C) unpaired sequences. (D) The bar plot shows the mean accuracy and standard error of the pair classification task. Metrics on classification tasks are mean and standard error, with the highest values bolded and the second highest values underlined.

IgBERT and IgT5 were trained with an approach closest to the curriculum strategy – these models were pre-trained using unpaired sequences and finetuned with a mix of unpaired and paired sequences. Despite this similar training approach and the large size of IgT5 (3B parameters), these models tend to perform poorly on specificity classification tasks. This suggests that model architecture (particularly rotary embeddings) and a larger emphasis on paired sequences is critical to high performance on these tasks.

## DISCUSSION

Recent studies on AbLMs have shown that the benefits of training on natively paired sequences can be sufficient to overcome a significant disadvantage in training data scale compared to unpaired sequences. The high cost and effort required to recover natively paired antibody sequences has sparked interest in methods that maximize the training value of these limited datasets, including supplementing paired sequences with unpaired data. Theoretically, this mixed training approach should still allow the model to learn critical cross-chain features while leveraging the greater diversity in much larger unpaired sequence datasets. However, it is not clear how best to combine these two data types to maximize training value.

We first performed a systematic analysis of model architectures and data formatting methods to understand their effect on mixed model training. The most impactful change was the use of rotary embeddings, which delivered a surprisingly large improvement in mixed-data model performance. Compared to absolute PE, RoPE greatly reduced the loss of homogeneous models (paired-only or unpaired-only) when tested with the heterologous data type. This is strong evidence that mixed-data models should use rotary embeddings whenever possible. In addition, we noted that mixed-data models using a single chain separator token (whether that be <cls> or <sep>) performed best, with the <cls> token showing a slight performance benefit in homogeneous models as well.

Next, we introduced a curriculum learning approach to training AbLMs, which allows for a more gradual transition from unpaired to paired sequences. Including both data types throughout the entire training course, with an increased emphasis on paired sequences toward the end of training, resulted in the best final model performance. We also showed that adjusting the parameters of the unpaired probability curve can differentially affect model performance on unpaired or paired sequences, meaning the curriculum approach can be tuned for specific performance objectives. Generally, optimizing for unpaired loss resulted in larger overall performance gains on downstream tasks, perhaps because unpaired sequences comprise the bulk of the training data.

We expected that one of the primary performance-enhancing benefits of supplementing paired training data with unpaired sequences would be the ability to train larger models for longer without overfitting. CurrAb, a 650M parameter curriculum model, validated this assumption by outperforming existing models on various classification tasks including native pairing and antigen specificity. The magnitude of improvement over IgBERT and IgT5 was surprising, particularly given that these models were finetuned with a mixture of paired and unpaired sequences. While this strategy is similar to our curriculum training approach in that the final training stages do not focus exclusively on a single data type, we note two differences that may underlie this performance disparity. First, it is possible that exposure to paired sequences only during finetuning is insufficient for the model to learn cross-chain features fully; instead, paired sequences may need to be introduced at the earliest stages of training to capture their full training value. Second, architectural optimizations such as rotary position embeddings, which are lacking in IgBERT and IgT5, may yield substantial performance improvements on downstream classification tasks, particularly those that involve paired antibody sequences.

More broadly, these findings emphasize the importance of generating larger datasets of paired antibody sequences, as paired-only models approach the performance of mixed models trained with substantially more data. In other words, the marginal training value of new data appears to strongly favor paired antibody sequences, despite the high cost of generating natively paired antibody sequences. It may also be useful to apply additional training methods that focus learning on the CDR regions, such as preferential masking^16^ and focal loss^3,17^, to use both the paired and unpaired data more efficiently. Beyond AbLMs, the ability of curriculum-based training approaches to prevent catastrophic forgetting is likely to be beneficial for a variety of biological models for which cross-domain expertise is important. For example, a curriculum model trained with a combination of proteins and antibodies could perform as well as a specialized antibody sequence or structure model while retaining an expert-level understanding of general proteins. Such a model could be very well-suited for tasks like antibody-antigen docking.

In summary, we report three important findings. First, RoPE significantly impacts the ability of AbLMs to generalize across unpaired and paired data, highlighting positional embeddings as a key but underappreciated component of mixed-data biological language models. Second, curriculum AbLMs typically perform best when optimized for unpaired performance, but the most important factor for final model performance is ensuring a sufficient emphasis on paired sequences throughout the entire training course. Finally, curriculum learning effectively mitigates catastrophic forgetting, making it a useful strategy for training biological models that exhibit uniformly high performance across multiple domains.

## METHODS

### Training Data

Sequence data was downloaded from the OAS^18^ on September 12th, 2024. In addition to sequences from the OAS, the paired dataset was supplemented with an internally generated dataset of ∼400k paired sequences. Filtering and clustering were performed as described in AntiRef^19^, with additional filtering for sequences containing ‘nan’ characters. The dataset clustered at 90% was chosen for both datasets, resulting in 151,764,423 unpaired sequences and 1,717,423 paired sequences in the full datasets. The final 650M-parameter model, CurrAb, was trained on the full dataset and used 96% of the data for training, with the remaining 4% left out for the evaluation and test sets.

For compute efficiency, initial tests in Figures 2-4 were trained with a downsampled dataset (20% of the full dataset), resulting in 30,352,885 unpaired sequences and 343,485 paired sequences. These datasets were similarly split to use 96% of the sequences for training and the remaining 4% for the evaluation and test sets. Test sets used for CE loss in Figures 2, 3 and 4 were the paired test set and a sample of ∼10k sequences from the unpaired test set (to match the number of sequences in the paired test set).

For the specificity classification tasks, two datasets were generated. CoV-specific sequences were downloaded from CoV-AbDab^20^ on November 11th, 2024 and an internal donor L1236. Flu-specific sequences were obtained from Wang et al.^21^ and filtered for paired sequences only. Healthy donor sequences were obtained from memory B-cell sequences from the Jaffe et al.^22^ and Phad et al.^23^ datasets and filtered to exclude any sequences in the train and evaluation datasets to prevent data leakage for both model sizes. For the HD-Flu-CoV classifications, the datasets were clustered by class at 99%, then combined with an equal number of sequences in each class, resulting in a final dataset with 4,398 sequences. For the HD-CoV classifications, the datasets were clustered by class at 95%, then combined with an equal number of sequences in each class, resulting in a final dataset with 27,442 sequences. Both classification datasets were split for 5-fold cross-validation with stratification. The same datasets were used for the unpaired versions of these tasks, but only the heavy chain sequence was provided.

For the pair classification task, the paired sequences from the full-data test set were filtered to exclude any sequences in the downscaled train and evaluation datasets, to prevent data leakage for both model sizes. This resulted in 41,564 paired sequences, which are comprised of both naive and memory B-cell sequences. The dataset was processed as described in Ng and Briney 2024^16^, including a split for 5-fold cross-validation with stratification.

### Curriculum Implementation

To enable training of different types of mixed models (constant and curriculum) modifications were made to the HuggingFace^24^ trainer using a trainer callback and by subclassing the PyTorch^25^ dataset class. This facilitated the separation of unpaired and paired datasets (for logging and evaluation purposes) and allowed for tracking and updating the unpaired probability function throughout training. The unpaired probability function determined what percentage of sequences should be sampled from the unpaired dataset at each step during training. The unpaired probability *P*(*t*) for the curriculum models is represented with the following equation:

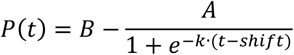

where *t* equals the current step divided by the total train steps. The values of *A* (height of the curve) and *B* (the vertical shift, aka upper bound of the curve) were modified to adjust the range of the probabilities, the *k* value was modified to adjust the slope of the sigmoid function, and the *shift* values were calculated to ensure that the total unpaired percentage (ie. 62.5%) was achieved. Refer to ***Table S6*** for the specific values used for each curriculum model.

### Model Pre-Training

All models used a slightly modified ESM-2 architecture model. This is an encoder-only architecture that produces embeddings useful for downstream tasks such as specificity classification, pair classification, and structure prediction. Models were trained with an MLM objective. This means that 15% of the input sequence was selected for prediction and of these, 80% were replaced with a <mask> token, 10% were replaced with a random token from the vocabulary, and 10% were left unchanged.

All models used a slightly modified ESM-2 vocabulary of 33 tokens, with an added <sep> token. Based on the separator tests presented in ***Table 1***, the large-scale models were trained with a <cls> separator. Sequences were preprocessed to place the separator between the heavy and light chains of the paired sequences, at the end of unpaired heavy chains, and at the beginning of unpaired light chains. Inputs were padded to 320 tokens, to accommodate the longest paired sequence.

For initial tests, we used a modified 55M-parameter ESM-2 architecture model, with 5 layers, 20 attention heads per layer, a hidden size of 960, and an intermediate size of 3840. After initial embedding tests in ***Fig 2***, RoPE was used for all subsequent models. Models were trained for 100k steps with a total batch size of 512. On 4 L40S graphics processing units (GPUs) using DeepSpeed ZeRO Stage 1^26^ via the Accelerate^27^ library, this equates to ∼11 hours per model. The peak learning rate was 4e-4, with a linear warmup for the first 6,000 steps followed by a linear decay. The only exception to this was the LR tests in Fig 3E-F, which tested WSD and SDGR LR schedules. The ideal parameters for WSD and SGDR were chosen based on Hägele et al 2024.^28^

The final curriculum model used the 650M parameter ESM-2 architecture, with RoPE and used the unique <cls> token as the separator token in the paired and unpaired sequences. The curriculum parameters were chosen to balance unpaired and paired performance – the max0.7 unpaired probability schedule was selected to benefit unpaired performance while k=20 was chosen to benefit paired performance. The model was trained for 500k steps with a total batch size of 512, equating to ∼6 days on 8 A100 GPUs using DeepSpeed ZeRO Stage 1. The peak learning rate was reduced to 1e-4 to assist with model convergence, with a linear warmup for the first 30,000 steps followed by a linear decay.

Models were logged using Weights & Biases (wandb). Data from wandb was used to produce the unpaired probability, LR schedule, and evaluation loss curves.

### Model Evaluation & Testing

CE loss and accuracy calculations on the eval and test sets were performed using the HuggingFace trainers evaluate function, with a custom compute_metrics function provided to ensure the metrics exclude the separator tokens from calculations. Accuracy was calculated using scikit-learn and CE loss was calculated using PyTorch.

CDRH3 accuracy was calculated by masking and predicting each position in the CDRH3 region individually, then calculating the accuracy of those predictions by averaging.

### Classification Tasks

For all classification tasks, models were finetuned with a standard sequence classification head. For all models we trained and models hosted on HuggingFace (IgBERT, IgT5, and AntiBERTa2), the base models were loaded with the default sequence classification head for that model architecture. For AbLang2, which is only available as a pip package, the sequence classification head was added manually and modeled after the ESM-2 classification head.

For specificity classification tasks, all models had a warmup ratio of 0.1 and a peak learning rate of 5e-5 followed by a linear decay. For binary classification (HD vs CoV), models were trained for 10 epochs and a total batch size of 32 per step. For 3-class classification (HD vs Flu vs CoV), models were trained for 5 epochs and a total batch size of 8 per step.

For pair classification tasks, models were finetuned following the training schedule presented in Ng and Briney 2024^16^. Models were trained for 50 epochs, with a warmup ratio of 0.1, a peak learning rate of 1e-5, and a total batch size of 256 per step.

Metrics used to evaluate the classifiers were: accuracy, F1, area under the receiver operating characteristic curve (**AUC**), area under the precision-recall curve (**AUPR**), and Matthews correlation coefficient (**MCC**). Metrics on the test set were averaged across cross-validation runs and standard error was calculated. Scikit-learn was used to calculate these metrics.

## Code and Data Availability

The code used for model training, evaluation, and figure generation is available on GitHub (github.com/brineylab/curriculum-paper). The training data and model weights for CurrAb are available on Zenodo (doi.org/10.5281/zenodo.14661302) and CurrAb is also available to use through Hugging Face (huggingface.co/brineylab/CurrAb).

## AUTHOR CONTRIBUTIONS

Conceptualization: SB, BB

Model training and evaluation: SB

Manuscript preparation and revisions: SB, BB

## SUPPLEMENTARY INFORMATION

**Table S1.** CE Loss and accuracy for separator token tests. Mixed, paired-only, and unpaired-only models were trained with 5 different separators. The separators tested were: no separator, <cls>, <cls><cls>, <sep>, and <sep><sep> where <cls> is the BOS token and <sep> is a unique separator token. Separators are placed between chains in paired sequences and unpaired sequences based on the chain (end of the heavy chains and the beginning of the light chains). Models were assessed on paired and unpaired test datasets, each containing ∼10k sequences.

| Separator | *Mixed Models* |  |  |  | *Paired Models* |  | *Unpaired Models* |  |
| --- | --- | --- | --- | --- | --- | --- | --- | --- |
|  | <u>Paired Data</u> |  | <u>Unpaired Data</u> |  | CE Loss | Accuracy | CE Loss | Accuracy |
|  | CE Loss | Accuracy | CE Loss | Accuracy |  |  |  |  |
| None | 0.1835 | **0.9510** | 0.3571 | **0.9133** | 0.1856 | **0.9507** | 0.3404 | 0.9177 |
| <cls> | **0.1828** | <u>0.9509</u> | **0.3561** | **0.9133** | **0.1851** | <u>0.9506</u> | <u>0.3389</u> | <u>0.9179</u> |
| <sep> | <u>0.1831</u> | <u>0.9509</u> | <u>0.3565</u> | <u>0.9131</u> | <u>0.1852</u> | <u>0.9506</u> | **0.3386** | **0.9180** |
| <cls><cls> | 0.1835 | 0.9508 | 0.3601 | 0.9125 | 0.1854 | 0.9506 | 0.3423 | 0.9175 |
| <sep><sep> | 0.1833 | 0.9508 | 0.3604 | 0.9124 | 0.1856 | 0.9505 | 0.3427 | 0.9174 |

**Table S2.** CE loss and accuracy for different unpaired percentages. Mixed models were trained with increasing percentages of unpaired data. Models were assessed on paired and unpaired test datasets, each containing ∼10k sequences.

| Unpaired % | Paired |  | Unpaired |  |
| --- | --- | --- | --- | --- |
|  | CE Loss | Accuracy | CE Loss | Accuracy |
| 37.5 | **0.1831** | 0.9508 | 0.3635 | 0.9111 |
| 50 | **0.1831** | **0.9509** | 0.3565 | 0.9131 |
| 62.5 | 0.1833 | **0.9509** | 0.3508 | 0.9148 |
| 75 | 0.1839 | 0.9508 | **0.3463** | **0.9164** |

**Table S3.**
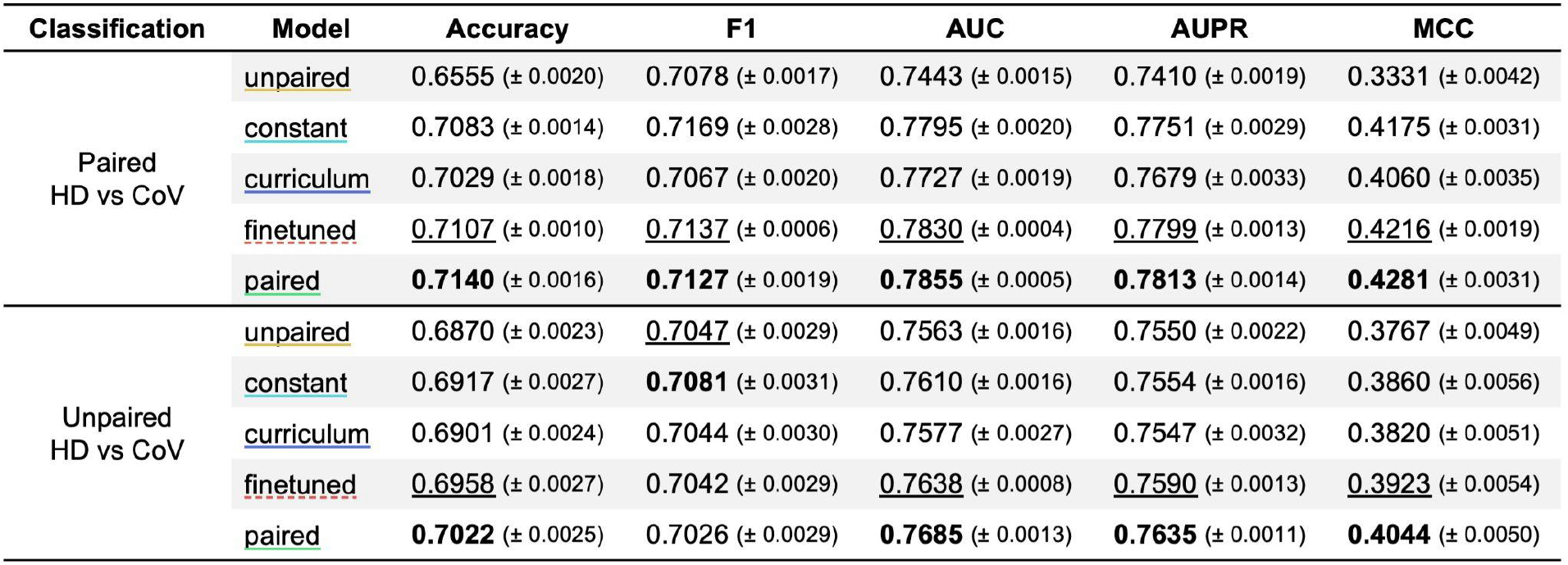
Additional classification tasks to compare mixed model training methods. Results for ‘HD vs CoV’ specificity classification task with paired and unpaired sequences. Metrics on classification tasks are mean and standard error, with the highest values bolded and the second highest values underlined

| Classification | Model | Accuracy | F1 | AUC | AUPR | MCC |
| --- | --- | --- | --- | --- | --- | --- |
| Paired<br>HD vs CoV | <u>unpaired</u> | 0.6555 (± 0.0020) | 0.7078 (± 0.0017) | 0.7443 (± 0.0015) | 0.7410 (± 0.0019) | 0.3331 (± 0.0042) |
| | constant | 0.7083 (± 0.0014) | 0.7169 (± 0.0028) | 0.7795 (± 0.0020) | 0.7751 (± 0.0029) | 0.4175 (± 0.0031) |
| | <u>curriculum</u> | 0.7029 (± 0.0018) | 0.7067 (± 0.0020) | 0.7727 (± 0.0019) | 0.7679 (± 0.0033) | 0.4060 (± 0.0035) |
| | <u>finetuned</u> | <u>0.7107</u> (± 0.0010) | <u>0.7137</u> (± 0.0006) | <u>0.7830</u> (± 0.0004) | <u>0.7799</u> (± 0.0013) | <u>0.4216</u> (± 0.0019) |
| | <u>paired</u> | **0.7140** (± 0.0016) | **0.7127** (± 0.0019) | **0.7855** (± 0.0005) | **0.7813** (± 0.0014) | **0.4281** (± 0.0031) |
| Unpaired<br>HD vs CoV | <u>unpaired</u> | 0.6870 (± 0.0023) | <u>0.7047</u> (± 0.0029) | 0.7563 (± 0.0016) | 0.7550 (± 0.0022) | 0.3767 (± 0.0049) |
| | constant | 0.6917 (± 0.0027) | **0.7081** (± 0.0031) | 0.7610 (± 0.0016) | 0.7554 (± 0.0016) | 0.3860 (± 0.0056) |
| | <u>curriculum</u> | 0.6901 (± 0.0024) | 0.7044 (± 0.0030) | 0.7577 (± 0.0027) | 0.7547 (± 0.0032) | 0.3820 (± 0.0051) |
| | <u>finetuned</u> | <u>0.6958</u> (± 0.0027) | 0.7042 (± 0.0029) | <u>0.7638</u> (± 0.0008) | <u>0.7590</u> (± 0.0013) | <u>0.3923</u> (± 0.0054) |
| | <u>paired</u> | **0.7022** (± 0.0025) | 0.7026 (± 0.0029) | **0.7685** (± 0.0013) | **0.7635** (± 0.0011) | **0.4044** (± 0.0050) |

**Table S4.** Classification tasks on 650M-parameter paired-only and curriculum models. Results specificity classification tasks (HD vs CoV’ and ‘HD vs Flu vs CoV’) with paired and unpaired sequences and the pair classification task. Metrics on classification tasks are mean and standard error, with the highest values bolded.

| Classification | Dataset | Model | Accuracy | F1 | AUC | AUPR | MCC |
| --- | --- | --- | --- | --- | --- | --- | --- |
| HD vs CoV | Paired | <u>Paired-650M</u> | 0.7450 (± 0.0010) | 0.7487 (± 0.0011) | 0.8242 (± 0.0008) | 0.8233 (± 0.0016) | 0.4901 (± 0.0020) |
| | | <u>CurrAb</u> | **0.7626** (± 0.0023) | **0.7669** (± 0.0016) | **0.8452** (± 0.0015) | **0.8381** (± 0.0023) | **0.5255** (± 0.0045) |
| | Unpaired | <u>Paired-650M</u> | 0.7180 (± 0.0008) | 0.7281 (± 0.0001) | 0.7953 (± 0.0015) | 0.7964 (± 0.0014) | 0.4373 (± 0.0017) |
| | | <u>CurrAb</u> | **0.7218** (± 0.0014) | **0.7316** (± 0.0012) | **0.8011** (± 0.0010) | **0.7992** (± 0.0014) | **0.4448** (± 0.0027) |
| HD vs Flu<br>vs CoV | Paired | <u>Paired-650M</u> | 0.6567 (± 0.0015) | 0.6571 (± 0.0014) | - | - | 0.4853 (± 0.0022) |
| | | <u>CurrAb</u> | **0.6796** (± 0.0051) | **0.6801** (± 0.0051) | - | - | **0.5200** (± 0.0076) |
| | Unpaired | <u>Paired-650M</u> | 0.6260 (± 0.0057) | 0.6266 (± 0.0058) | - | - | 0.4397 (± 0.0088) |
| | | <u>CurrAb</u> | **0.6401** (± 0.0052) | **0.6408** (± 0.0053) | - | - | **0.4612** (± 0.0077) |
| Native vs<br>Random Pairs | - | <u>Paired-650M</u> | 0.6373 (± 0.0017) | **0.5618** (± 0.0024) | 0.6920 (± 0.0011) | 0.7189 (± 0.0014) | 0.2926 (± 0.0036) |
| | | <u>CurrAb</u> | **0.6428** (± 0.0008) | 0.5462 (± 0.0029) | **0.6985** (± 0.0017) | **0.7289** (± 0.0016) | **0.3156** (± 0.0011) |

**Table S5.**
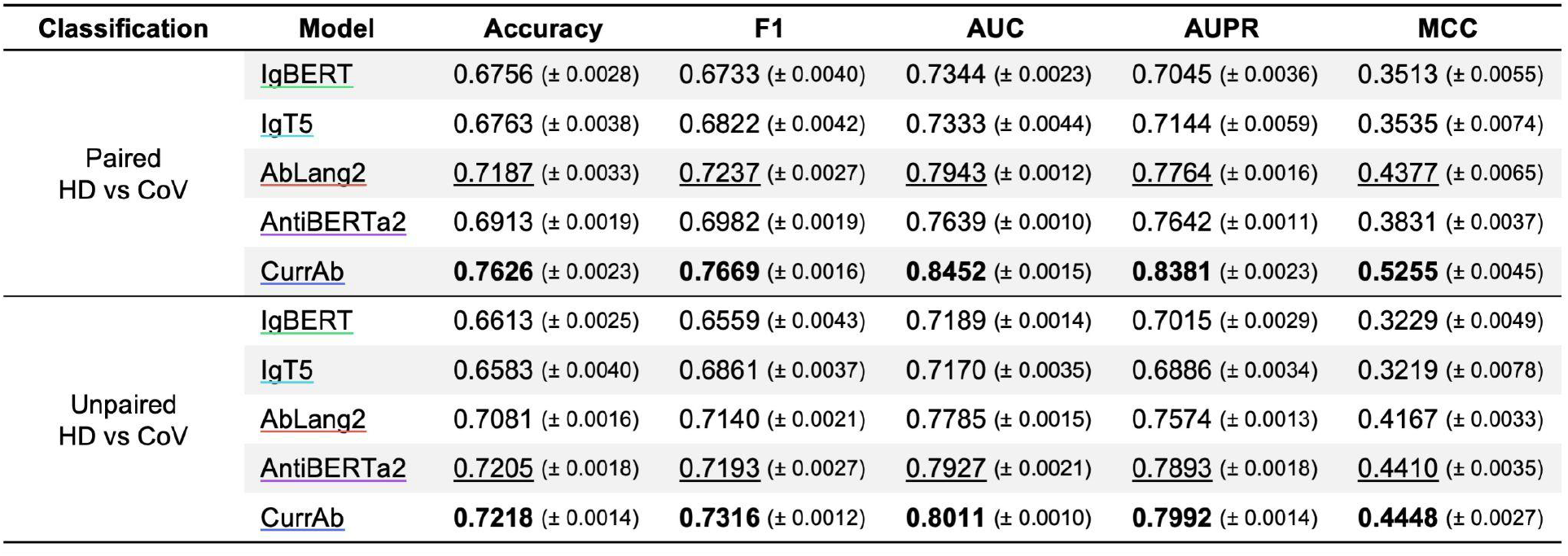
Additional classification tasks to compare large-scale models. Results for ‘HD vs CoV’ specificity classification task with paired and unpaired sequences. Metrics on classification tasks are mean and standard error, with the highest values bolded and the second highest values underlined.

**Table S6.**
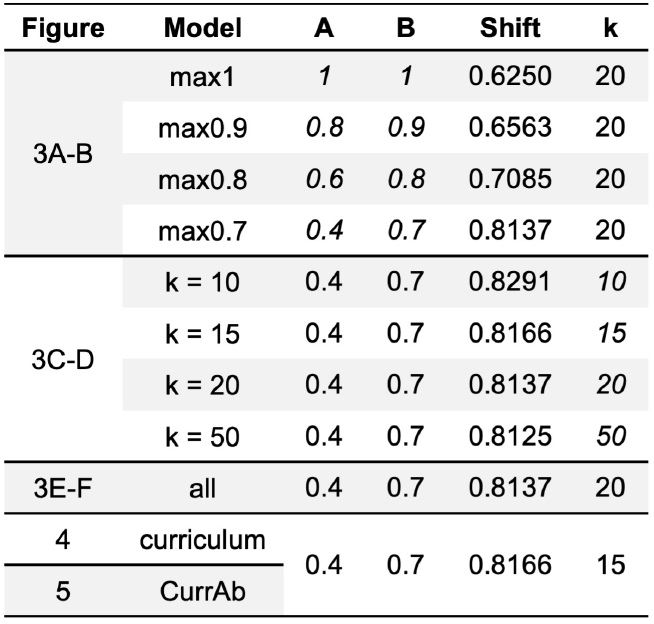
Unpaired probability equation values for curriculum models. Corresponding figure, model, and equation values (A, B, shift, and k) are listed for each model. Italics represent the values being tested in the given figure.

